# Potential pitfalls in estimating viral load heritability

**DOI:** 10.1101/046797

**Authors:** Gabriel E. Leventhal, Sebastian Bonhoeffer

## Abstract

In HIV patients, the set-point viral load (SPVL) is the most widely used predictor of disease severity. Yet SPVL varies over several orders of magnitude between patients. The heritability of SPVL quantifies how much of the variation in SPVL is due to transmissible viral genetics. There is currently no clear consensus on the value of SPVL heritability, as multiple studies have reported apparently discrepant estimates. Here we illustrate that the discrepancies in estimates are most likely due to differences in the estimation methods, rather than the study populations. Importantly, phylogenetic estimates run the risk of being strongly confounded by unrealistic model assumptions. Care must be taken when interpreting and comparing the different estimates to each other.

## Searching for set-point viral load heritability in HIV infection

During the asymptomatic phase of HIV infection, the viral load within a patient fluctuates around a relatively stable value known as the set-point viral load. Set-point viral load (SPVL) has proven to be highly relevant, as untreated patients with higher SPVL tend to progress to AIDS faster than those with low SPVL, and consequentially SPVL is one of the most widely used predictors of disease severity [1, 2]. What is particularly striking, however, is the amount of variation in SPVL between patients: SPVL varies over several orders of magnitude between patients [1, 3–5]. Understanding the source of this wide variation in SPVL in the patient population is key to understanding HIV pathogenicity and why certain patients progress to AIDS rather quickly, while others progress much more slowly or not at all.

The potential contributions to SPVL variation can generally be split into four categories, determined by:

i. the specific genetic viral variant that infects a host;
ii. the host genetics;
iii. any interactions between host and viral genetics;
iv. other extrinsic factors, both independent of and interacting with host and viral genetics.

From a virus-centric view, we can group contributions (ii)-(iv) together, and hence the question becomes to what degree the variation in HIV genetics explains the variation in SPVL in the patient population, and by proxy how much viral genetics control the variation HIV pathogenicity.

New HIV infections can occur after a virus is transmitted from a donor host to a recipient host. Thus, from an evolutionary point of view, viral genetic information is passed on and conserved from one infection to the next, whereas donor and recipient host genetics are typically unrelated. In evolutionary theory, the concept of heritability quantifies precisely the question at hand [6]: How much of the observed variation in a trait is explained by variation in the genetics that are passed on to the next generation? A heritability of 100% means that all of the trait variation is explained by transmissible genetic information, while a heritability of 0% means that transmissible genetics explain none of the trait variation. A number of recent publications have reported heritability estimates based on different methods that range from as low as 5% to as high as 50% [7–14]. These studies used partly different methods to estimate heritability, depending on the available data. The crucial question thus arises as to whether these discrepant estimates reflect true differences in the study populations, or whether the differences may rather be artifacts of the estimation method used.

Every statistical model comes with a set of assumptions that must be fulfilled in order for the results to be meaningful. Here, we aim to illustrate how some of these estimates of heritability might be confounded by model assumptions that do not completely conform to the dynamics of HIV transmission. Nevertheless, even when the assumptions are fulfilled, the choice of estimation method alone can potentially lead to differences in the heritability estimates.

## Estimating the heritability of a viral trait

The concept of heritability was initially developed to measure the degree of correspondence between the trait value in an offspring and the trait value of its parents, and can be traced back to Galton [15], though the first use of the term “heritability” remains elusive [16]. Most of the credit for developing the methods to estimate heritability go back to Fisher [17] and Wright [18], although the common contemporary use corresponds to “narrow sense heritability” as defined by Lush [19]: Heritability is the ratio of additive genetic variance, 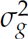, and the sum of additive genetic and environmental variance, 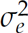, in a population,

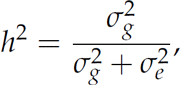

where the total phenotypic variance, 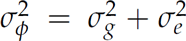, is the sum of all contributions to variance. Heritability can then be estimated from data of related individuals (e.g. parents and their offspring) using regression methods [6, 17, 18]. For the case of SPVL in HIV, 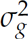 corresponds to the transmissible contribution (i) and 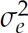 comprises the non-transmissible contributions (ii)–(iv).

Analogous to these methods that were developed for sexually reproducing individuals, the trait variance of clonally reproducing organisms (such as viruses) can be partitioned into variance components. In viral infections, only the virus genotype is passed on from one infection to the next, and thus the heritability of a viral trait as defined by *h*^2^ can be used to quantify the relative influence of the variation in transmissible viral factors to the trait of interest in a population [8, 20]. While the general framework of partitioning trait variance is the same in sexual and clonal populations, the underlying processes that generated the observed variance are different. Thus, although the methods to estimate heritability in sexual populations can generally be directly applied to viral heritability, care must be taken when evaluating the applicability of the methods.

Furthermore, whenever heritability is estimated, it is also important to remember that even though *h*^2^ can be used to characterize the propensity of virus genetics to influence a trait, the actual measured heritability depends on both the expressed variation of this trait in a given population, as well as the variation in other factors that influence the trait. Therefore, *h*^2^ strongly depends on the population it is measured in, and can consequently vary between study populations. This does not invalidate the usefulness of measuring *h*^2^, but rather means that predictions on the basis of *h*^2^ are primarily valid in the study population at hand and care must be taken when using *h*^2^ for predictions in other populations.

### Parent-offspring regression

Parent-offspring regression is tightly linked to the original definition of heritability and is thus the most straightforward estimation method [17, 18]. The idea is to compare trait values in parents to trait values in their respective offspring, or in the case of viral infections compare traits from donors and recipients within transmission pairs. The applicability of “donor-recipient” regression (DR) in the context of viral heritability has been recently reviewed in Fraser *et al.* [8]. In short, DR assumes that viral traits are imperfectly passed on from one infection to the next and determines the predictive ability of SPVL in a donor of SPVL in the recipient. Differences in SPVL between donors and recipients must then either be due to non-transmissible contributions of type (ii)-(iv), or due to genetic changes that accumulate as a result of intra-host evolution or genetic bottlenecks at transmission [20]. Viral heritability of SPVL is then the regression slope of recipient SPVL on donor SPVL, akin to parent-offspring regression [6].

The biggest difficulty in estimating heritability using donor-recipient regression lies in observing direct transmission between pairs. Currently, the best source of such transmission pairs are discordant couple study cohorts [8]. But even then, there is the possibility that infection of the seronegative partner occurred from an unknown third-party. Furthermore, such studies are restricted to specific study populations that may not be representative of the general global HIV epidemic. Non-independence between different donor-recipient pairs such as a mutual infector of the donors may further distort estimates.

### Phylogenetic methods

To overcome the shortage of appropriate study populations, considerable hope has been placed in estimating heritability from much more easily available sequence information using phylogenetic methods. Various phylogenetic methods have previously been employed to estimate the heritability of SPVL [7, 9, 21]. All of these methods have been developed in the context of ecological species and have yet to be validated for use on viral populations. The methods used include non-parametric estimates of phylogenetic signal [22], as well parametric regression methods such as independent contrasts [23], the phylogenetic mixed model [24], and Pagel’s *λ* [25]. Non-parametric methods can be indicative of the level of signal in a dataset, but are generally less useful when the goal is a quantitative estimate of heritability. The parametric regression methods are in essence an extension of donor-recipient regression that can be applied to datasets where the relation between individuals goes beyond parents and offspring or direct transmission pairs. A consequence of this extension is that a process model is required that links the observed genetic differences to the number of transmission events and the time subjected to intra-host evolution. All of the above methods assume that the genetic differences accumulate neutrally through time, irrespective of the number of transmission events or selection on the trait of interest, which generally is not the case for HIV. Hence, the reported estimates must therefore be interpreted together with such model assumptions. Despite being developed from different first principles, in certain cases Pagel’s *λ* is equivalent to the Phylogenetic Mixed Model (Box 1). In the following sections we thus only consider the Phylogenetic Mixed Model (PMM), which is, as shown in Box 1, the most generic and appropriate method for viral heritability estimation.

## Heritability estimation in simulated populations

In order to illustrate the effects of incorrect model assumptions about the evolutionary process underlying the phylogenetic methods, we compare the different estimation methods using mock data from simulations (Fig. 1). While simulated data have previously been used to validate the respective heritability estimates for SPVL in HIV [7, 9, 26], the simulation models differ between publications and generally do not adequately represent key aspects of the evolution of SPVL across infections. Here, we use a simple previously published model for HIV transmission across many generations in a Wright-Fisher population that accounts for transmission bottlenecks and intra-host evolution [20].

**Figure1:**
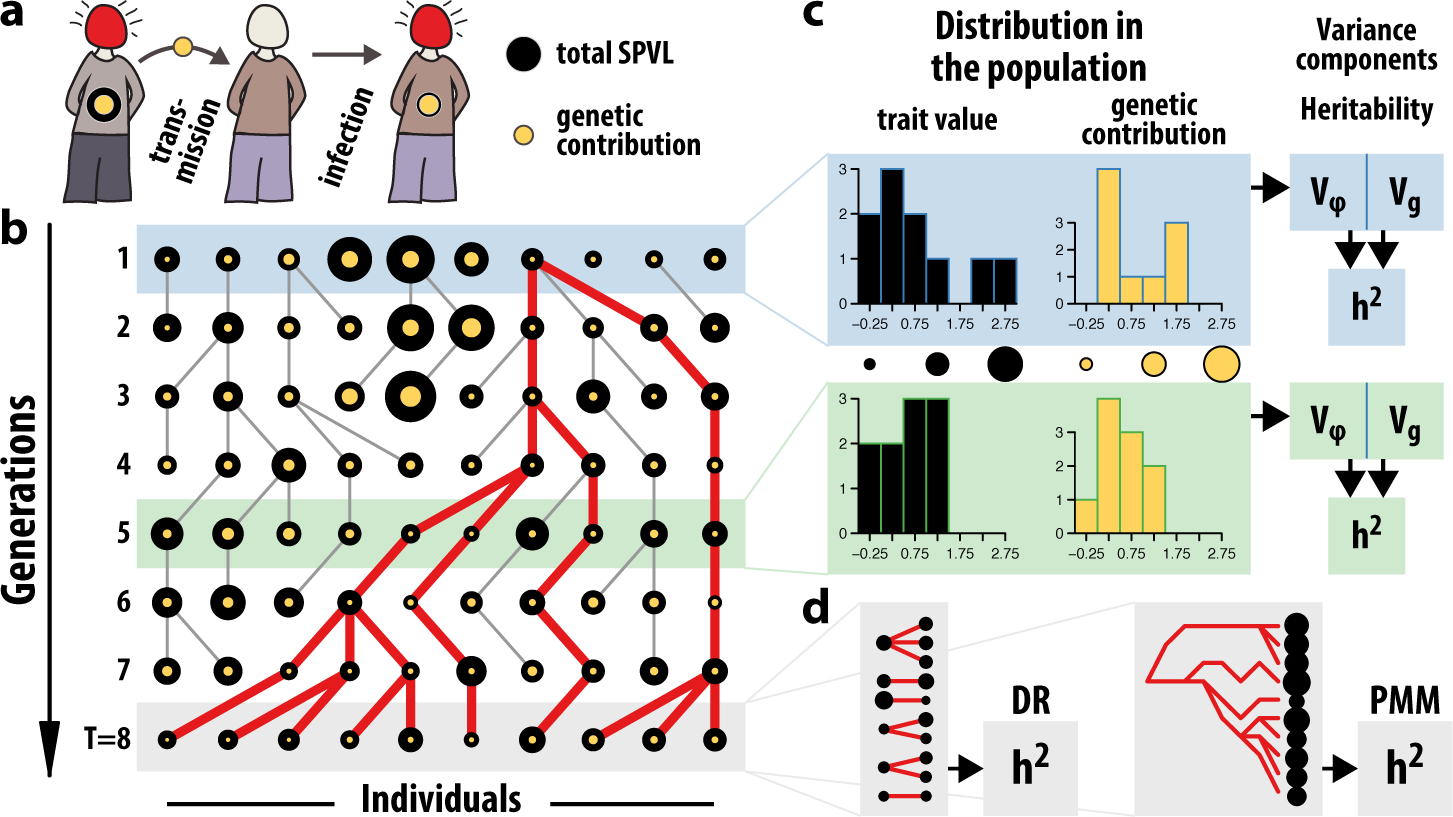
Estimating heritability in a simulated Wright-Fisher population. **(a)** At transmission, only the genetic contribution to SPVL is passed on from donor to recipient.Differences in host contribution result in differences in SPVL between donors and recipients with the same genetic contribution. **(b)** At every generation, parents are chosen from the previous generation. The circle size of each individual is representative of its trait value. The genetic component (yellow circles) is passed on to the offspring including potential genetic changes due to intra-host evolution. The red lines show the full ancestry of the individuals in the last generation. **(c)** The distribution of the trait in the population may change from generation to generation and is shown for two particular generations. The heritability in a population at a specific generation time is the genetic variance divided by the trait variance. **(d)** Alternatively, the heritability in the final generation can be estimated using either a donor-recipient regression or a phylogenetic method.

An individual’s log SPVL is assumed to be a linear combination of both a transmissible component, *g*, and a non-transmissible component, *e*, such that SPVL of individual *i* at generation *t* is *ϕ_i_*(*t*) = *g_i_*(*t*) + *e_i_*(*t*). For each individual *i* in the subsequent generation, a donor, *d*(*i*), is chosen from the current generation, either uniformly (drift) or based on their value of SPVL (selection). The genetic contribution of the donor is then transferred from to the recipient, 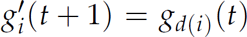, including an potential additional change due to intra-host evolution that follows a normal distribution with variance 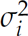. Because the population is finite, there is a generation *T* at which the whole population will have descended from the same individual (Fig. 1), and the phylogeny or pedigree of the population can be traced back to the common ancestor. Thus we can compare heritability estimates in this final generation obtained using DR regression or the PMM.

We first consider the case where SPVL does not influence the likelihood of transmission, and hence there is no selection on SPVL between hosts, since this is a key assumption underlying the PMM. Note that the distribution in the genetic contribution to SPVL may still change from generation to generation due to drift. The heritability estimated by donor-recipient regression is consistent with the fraction of variances in the final generation, 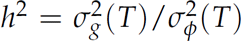 (Fig. 2a). Heritability estimates found using the PMM, however, do not estimate the ratio of variances, but rather the amount of genetic variance that would be expected from a drift process that has run for *T* generations, with an increment in variance per generation, 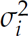, equal to that produced by intra-host evolution (Fig. 2b). The expected amount of genetic variance at generation *T* thus is 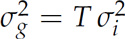, which is generally not equal to the variance estimate in the final generation. In addition to estimating differing quantities, PMM also strongly underperforms in in precision compared to DR regression when only a fraction of the population is sampled (Fig. 2c).

**Figure 2:**
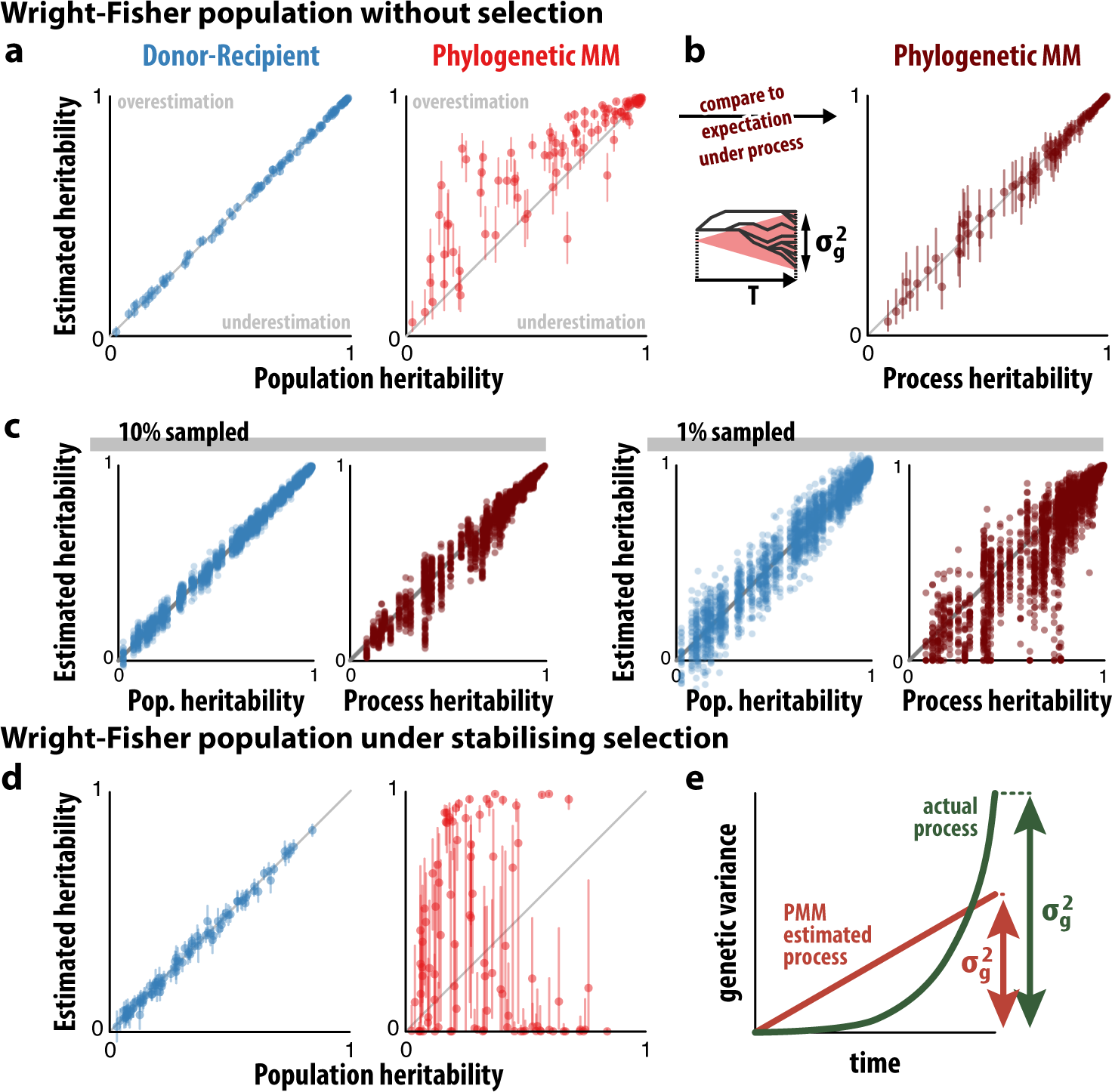
Heritability estimation using donor-recipient regression and the Phylogenetic Mixed Model. Wright-Fisher populations of 10000 individuals were simulated as described in Fig. 1. At each generation, the genetic contribution in recipients is chosen from a normal distribution centered on the value of the donor with variance 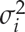 (see also [20]). **a.** Maximum likelihood heritability estimates of DR regression (blue) and the PMM (red) compared to population heritability as defined by the ratio of variances. Vertical lines indicate the profile likelihood confidence intervals. **b.** The PMM consistently estimates the process heritability based on the expected genetic variance at generation *T*, given a per-generation incremental increase of 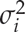 (dark red). **c.** Estimates from DR regression remain robust even at low sampling levels of 10% and 1%, compared to the PMM. DR and PMM estimates are compared to the population and process heritability, respectively. **d.** Heritability estimates using DR and PMM when SPVL influences the likelihood of transmission. **e.** The PMM estimates a linear variance generating process, which can be different from the actual process that generated the variance. For **a-c**, 100 random parameter sets were uniformly chosen from 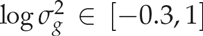, 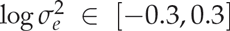, 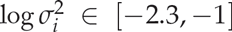. For **d**, 100 random parameter sets were chosen acording to the true transmission potential for HIV [5]: 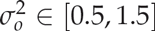, 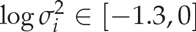, 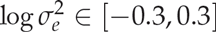, 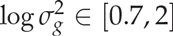.

Next we consider the case where SPVL influences the likelihood of transmission of an individual, as is known for HIV [5]. We compare both the PMM and DR to the population heritability, i.e. the fraction of variance components, as there is no equivalent process based interpretation of heritability for a non-neutral trait. Because DR regression only depends on a single transmission event, the estimate is robust to the selection of donors based on SPVL (Fig. 2d; see also [8]). Heritability estimates obtained using the PMM generally break down if the population evolved under selection on the trait of interest. This can lead to an unpredictable underestimation or overestimation bias, and also misleading narrow confidence intervals (Fig. 2d). Such overconfidence in biased estimates is typical of fitting parametric models. A certain parameter value might very clearly give the best fit of the underlying process to the data and hence that parameter value has narrow confidence intervals. However, the actual model process might still be a poor fit to the data. For a population evolving under selection on the trait of interest, this generally leads to an underestimation of the genetic contribution to variance, 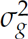, when using the PMM (Fig 2e). However, depending on the esimate of the overall variance, 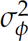, under the PMM (Box 1), this unpredictably leads to either overestimation or underestimation of heritability under the PMM (Fig 2d).

## Comparing donor-recipient and phylogenetic estimates

As we have illustrated, estimates from DR regression and the PMM cannot be immediately compared to each other. Donor-recipient regression estimates the heritability in the current population under the assumption that all individuals are independent. This corresponds to calculating the sample variance of the genetic contributions of all individuals in the populations, and dividing by the sample variance of the phenotype of all individuals in the population. It is therefore a measure of the sampled population at the current point in time, irrespective of the evolutionary history. In this sense, donor-recipient regression is robust to variations in the evolutionary model that generated the genetic variance in the population. The phylogenetic mixed model estimates the process heritability under the assumption that the genetic contribution to the variance in the population increased linearly through time, i.e. through drift. This corresponds to calculating the expected variance in a population that has evolved from a common ancestor at some time *T* in the past. This latter heritability is therefore a measure of the process that gave rise to the current population and thus depends on the evolutionary history of the population. Hence, a mismatch between the evolutionary process that is assumed in the PMM and the true process that occurred can strongly confound the estimates in an unpredictable way. Viral phylogenies reconstructed from real data are also mostly non-ultrametric and consequently the process time *T* has to be chosen arbitrarily, which can further affect heritability estimates (see Box 2).

Which interpretation of heritability we are interested in depends on how *h*^2^ is used for prediction. One common use of *h*^2^ is to determine the amount of genetic variation that can be potentially found. A large *h*^2^ increases the statistical power of GWAS studies to detect common genetic variants that influence a trait [27, 28]. Another common use of *h*^2^ is to predict the response to selection [6]. A classic result in population genetics relates the magnitude of change in phenotype in the subsequent generation to the product of the heritability and the strength of selection. The larger *h*^2^, the larger the effect of selection on a population. Both of these uses are only concerned with genetic variance in the current population, irrespective of its evolutionary history. Therefore, *h*^2^ estimates from DR studies are most appropriate. We can also make use of heritability estimates to understand the evolution of HIV. From one infection to the next, two main processes can contribute to genetic changes: First, intra-host evolution can result in the accumulation of genetic changes in the viral population that may then be transmitted. Second, transmission bottlenecks may select a minority variants from the viral population which is then transmitted. When the assumptions about the process are correct, then phylogenetic mixed model can estimate the rate at which such transmissible genetic changes take place.

## Concluding remarks and outlook

The discrepancy in reported estimates has lead to some confusion about the reliability and usefulness of estimating heritability of SPVL. On the one hand, estimates using DR regression in various study populations from Africa and Europe produce consistent estimates (Tab. 1). On the other hand, estimates based on phylogenetic trees suggest potentially large differences between studies. However, these latter phylogenetic estimates are difficult to compare, as only Hodcroft *et al.* [9] employ the PMM, while the remaining studies report Pagel’s *λ* and other measures [7, 21, 26], from which heritability cannot readily be derived for non-ultrametric trees (see Box 1). Furthermore, the simulation studies used to validate these other measures do not produce the actual heritability that results from explicitly accounting for the transmissible genetic contribution to SPVL as we do here.

**Table 1:**
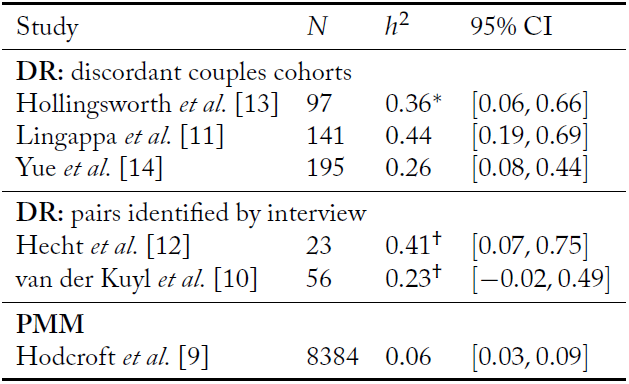
Reported heritability estimates. *Here we report the re-estimate of heritability based on the DR regression slope from [8]. ^†^The studies by Hecht *et al.* [12] and van der Kuyl *et al.* [10] reported Pearson correlation coefficients, rather than regression slopes. For these studies, we extracted the donor and recipient log_10_ SPVL values for the published figures (Fig. 2 in [12] and Fig. 1 in [10]), and recalculated the estimates of the regression slope. The precision of the estimates is thus dependent on the precision of the plotted points in the figures.

This leads us to speculate that most of the differences in heritability estimates come from differences in the estimation methods, rather than actual differences in the data. One must thus decide, which of the methods produces estimates that are more relevant to the research questions being asked. The increasing amount of available viral sequence data presents an intriguing opportunity to radically increase the number of data points. However, larger *n* does not guarantee better estimates if phylogenetic methods are used instead of donor-recipient regression. In our mock data, we see that phylogenetic methods potentially need at least one to two orders of magnitude more data to achieve similar confidence in the estimates. More importantly, the model assumptions that accompany the phylogenetic methods cannot be blindly ignored.

In HIV infection, a higher viral load results in a higher likelihood of transmission. The viral load in a patient varies over time, with the largest overall difference occurring between the relatively high viral load during the acute phase compared to the set-point viral load at the beginning of the chronic asymptomatic phase. Thus, whether this link between viral load and transmission rate translates into between-host selection on *set-point* viral load is not yet fully resolved and may depend on the relative contributions of onward transmission during the acute and the chronic phases. Yet, there is nevertheless evidence of a certain degree of balancing selection for optimal levels of SPVL in HIV on a between host level [5, 29, 30].

A proper estimation of heritability using phylogenetic trees would require extensions to the PMM that incorporate the relevant aspects of the true evolutionary process. For example, constant stabilizing selection on SPVL could be accounted for using an Ornstein-Uhlenbeck (OU) process [31]. This has previously been applied to study macro-evolution on ultrametric trees [32], and current efforts are underway to develop an OU-based method for non-ultrametric pathogen trees (personal communication, Venelin Mitov & Tanja Stadler, Apr 2016). As stabilising selection on SPVL has shown to occur [5, 29, 30], this may represent a major step forward in estimating heritability using phylogenetic methods. Yet in cases where the selective process is unknown or may have fluctuated over the evolutionary history, constructing an appropriate evolutionary model may nevertheless be challenging. Likelihood-free methods such as Approximate Bayesian Computation (ABC) have the advantage of being rather flexible in terms of the underlying evolutionary process. Such an approach has previously been applied to constrain the magnitude of selection and genetic change that would be compatible with the observed distribution of SPVL in patient populations without taking the phylogenetic relationships into account [20]. Incorporating phylogenetic information into such an ABC approach is currently not possible, as we still lack sufficiently large systematic HIV sequence data. Current large-scale collaborative projects such as BEEHIVE^(i)^ promise to fill this data gap.

Finally, throughout this paper we have assumed that the phylogenetic tree exactly represents the genetic relatedness of viral isolates from patients. In reality, phylogenetic trees are reconstructed from sequence data and are thus only estimates of the true phylogenetic tree. It is difficult to predict in which way errors in the tree might bias heritability estimates, but incorporating the uncertainty in the tree into the uncertainty in the heritability estimates is advisable and can readily be done in a Bayesian framework [21]. Overall, estimates from phylogenetic methods cannot be readily preferred over estimates from donor-recipient studies without properly scrutinizing the adequacy of the model assumptions that underlie the estimation method and taking into account additional noise and biases that may arise from the phylogenetic tree reconstruction.

## Acknowledgements

The authors thank Venelin Mitov, Tanja Stadler, Christophe Fraser, Samuel Alizon, Roland Regoes and Frederic Bertels for constructive discussions.

## Resources

(i) BEEHIVE: Bridging the Evolution and Epidemiology of HIV in Europe, https://erc.europa.eu/projects-and-results/erc-funded-projects/beehive. Last accessed: 12 Apr 2016.

## Glossary

Asymptomatic phase: HIV infections are generally split into three characteristic stages: (I) Primary infection/Acute phase; (II) Chronic asymptomatic phase; (III) AIDS phase.
Viral load: The density of virus in the blood of a patient. It is a proxy for the amount of virus in the rest of the body.
Set-point viral load: The viral load during the asymptomatic phase fluctuates around a remarkably stable level, the set-point viral load.
Genetic variance: Amount of variance in the trait (e.g. SPVL) that is due to differences in transmissible viral genetics.
Environmental variance: Amount of variance in the trait (e.g. SPVL) that is due to anything other then viral genetics.
Viral heritability: The fraction of phenotypic variance that is explained by transmissible genetic factors.
Donor-recipient regression: Method to estimate heritability by regressing the trait values in the recipients on the traits values of the donors.
Seronegative: Negative for HIV infection.
Serodiscordant couples: Sexual partnerships where only a single individual is infected.
Pedigree: Ancestral tree linking parents to their offspring.
Genetic bottlneck: Sudden decrease in population size where only a few genetic variants are selected. This leads to a drastic decrease in genetic variation in the population.
Genetic drift: Change in the genotype distribution over time due to the finite size of a population.

## Box 1: Equivalence between phylogenetic methods

For a single trait value evolving along a tree, it is straightforward to derive the maximum likelihood estimators for Pagel’s *λ* (PL) and the Phylogenetic Mixed Model (PMM) at a given level of heritability. To do this, both PL and the PMM assume that the sampled trait values are distributed according to a multivariate normal distribution with covariance matrix **Σ**(*h*^2^) or **Σ**(*λ*) for PMM and PL, respectively. Multivariate normal maximum likelihood estimators for both the mean, *μ*, and the overall variance, 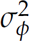, of a fixed covariance structure, **Σ**(*h*^2^), for a specific value of *h*^2^ are easily derived (Box 3). The maximum likelihood estimate of *h*^2^ can then be found by subsequently numerically maximizing the likelihood over values of *h*^2^ (or *λ* for PL). Despite following the same statistical framework, there are small differences in the derivation of **Σ** for PL and the PMM.

### The Phylogenetic Mixed Model

The idea of PMM is that the phenotype, *ϕ_i_*, of individual *i*, consists of a genetic, *g_i_*, environmental, *e_i_*, and a mean, *μ*, contribution,

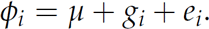

Under a linear statistical model, we can interpret the environmental contribution as a random error. We assume that the environmental contributions, ***e***, are drawn from the same distribution for all individuals,

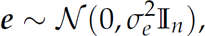

where 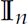 is the identity matrix with dimension *n*. We further assume that the genetic contributions, ***g***, are random samples from a multidimensional normal distribution with a covariance matrix equal to the genetic relatedness matrix **G** (Fig. I),

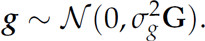

The phenotype distribution then is,

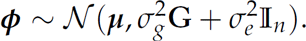

The variance of this normal distribution can also be written in terms of the total phenotypic variance, 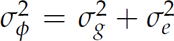, and the heritability, 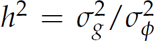, resulting in a final covariance matrix

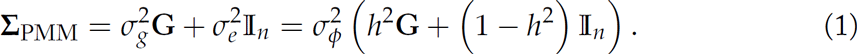

The entries *g_ij_* of **G** are proportional to the joint evolutionary time of the individuals *i* and *j* relative to some evolutionary origin *T* (Fig. I). If *a_ij_* is the time of the most recent common ancestor of individuals *i* and *j*, then *g_ij_* = 1 − *a_ij_*/*T*. The origin *T* is the time of the most recent common ancestor of all individuals, such that the smallest entry in **G** is 0 and the diagonal elements are 1.

### Pagel’s Lambda

The idea behind the PL method is to find an optimal rescaling parameter, *λ*, of the internal branches of the tree, such that the new ancestor times are *b_ij_* = *λa_ij_*, but the diagonal elements are unchanged, *b_ii_* = *a_ii_*. Under this rescaling, the covariance matrix is

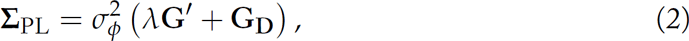

where **G***′* is equal to **G** with the diagonal removed, **G***′* = **G G_D_**.

### Comparing PL and PMM

A comparison between the PMM and PL becomes more clear by rewriting Eq. 1 in terms of **G***′* and **G**_*D*_,

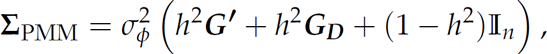

For ultrametric trees, all individuals are sampled at the same time *t_i_* = 0, and thus all the *g_ii_* = 1, and 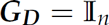. In this case,

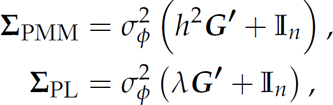

for which the equivalence between PL and PMM becomes evident by setting *h*^2^ = *λ*.

For non-ultrametric trees, not all *g_ii_* are the same and thus the two methods can no longer be made identical by simply setting *λ* = *h*^2^. The rescaling in PL only applies to the internal branches, which can be interpreted such that individuals that are sampled later and thus have larger entries in **G**_*D*_ also have larger environmental contributions. This, however, would only make sense if the environmental contribution increases linearly with time of sampling. A more sensible assumption is that the environmental contribution is independent of the time of sampling, and hence the PMM is the proper formulation of a neutrally evolving trait along a tree.

**Figure I:**
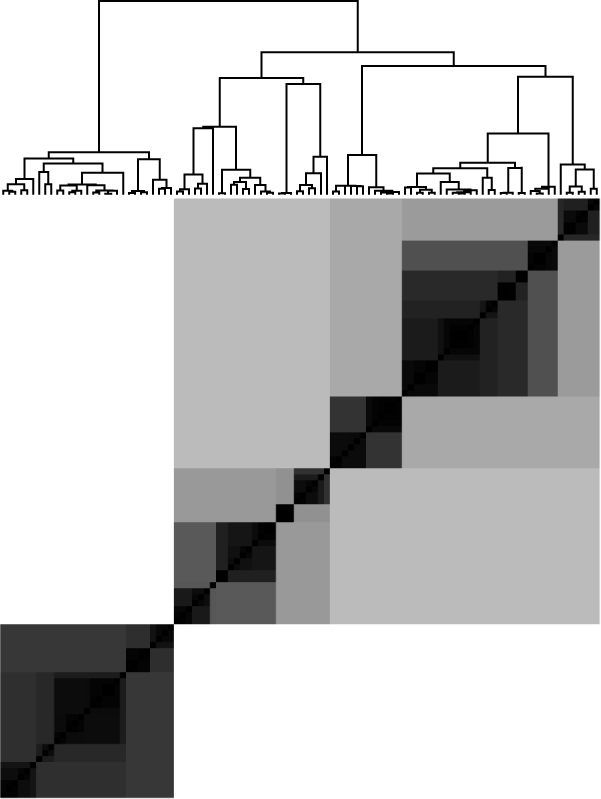
Graphical representation of the genetic relationship matrix for a Wright-Fisher population. The strength of the covariance between two individuals depends on the amount of shared ancestry in the tree. White entries corresponds to a covariance of 0 and black to a covariance of 1.

**Figure II:**
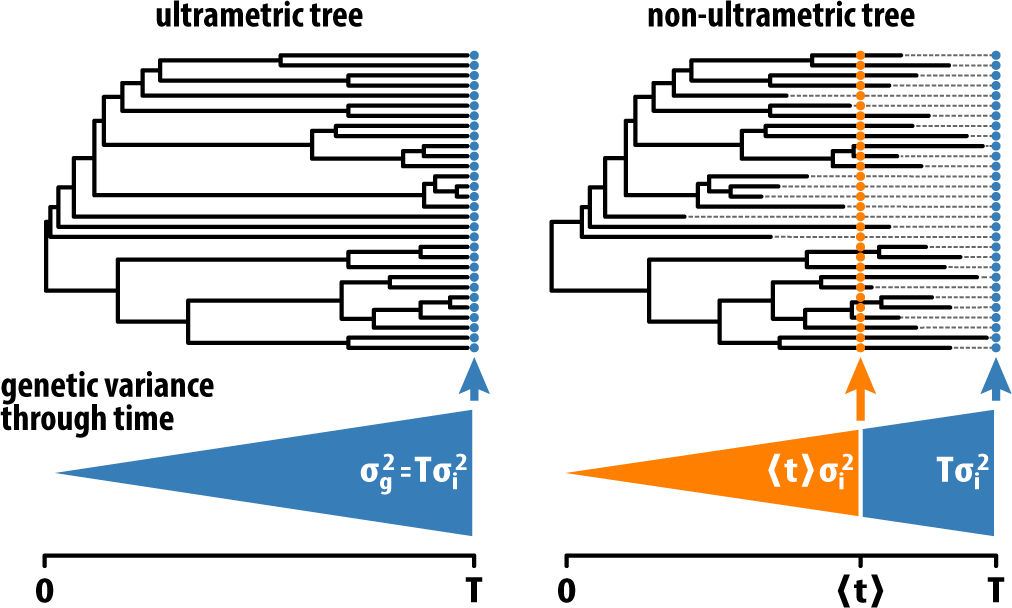
The PMM estimates a per-unit increase in genetic variance. For ultrametric trees, the estimated genetic varaince in the population is equal to the total height of the tree, *T*, times the estimated per-unit increase in genetic variance, 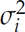. For non-ultrametric trees, the choice of the process time determines the amount of estimated genetic variance and hence the heritability. Common choices are the mean sampling time of all the tips, ⟨*t*⟩, or the total height of the tree, *T*.

## Box 2: Care must be taken when applying the PMM to non-ultrametric trees

An issue with applying the PMM to non-ultrametric trees is that the ancestry matrix, ***A***, needs to be rescaled by an arbitrary parameter. The entries, *a_ij_*, of the ancestry matrix represent the total time of shared ancestry of two individuals between the present and the root of the tree at some time *T* in the past. For ultrametric trees, all the tips of the tree align to the present, and hence the diagonal elements of ***A*** are all *T*. Thus, an obvious choice of rescaling is ***G*** = ***A***/*T*. The genetic variance component, 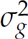, estimated using the PMM is then interpreted as the expected amount of genetic variance that accumulated during the time *T*. The per unit time increase of genetic variance, 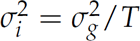, is the parameter of the underlying model of linear increase in variance in the PMM (Fig. II).

For non-ultrametric trees, the tips in the tree no longer all align to the present. Thus, the choice of normalization for ***A*** is less evident. We can write Eq. 1 from Box 1 using the ancestry matrix and 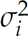 directly,

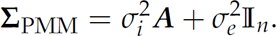

The mathematical framework to estimate variance components using the PMM thus remains valid for covariance matrices from non-ultrametric trees and can be used to estimate the variance component 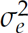 and the per unit time variance component 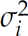. However, in order to estimate the heritability, 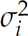 must be multiplied by an arbitrary time *τ*,

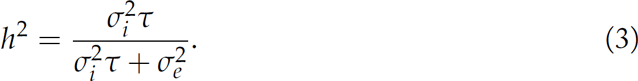

Equation 3 gives the heritability estimate for a population at time *τ* since the beginning of the process at the root of the tree. For ultrametric trees, choosing *τ* = *T* is the same as rescaling the ancestry matrix, ***A***, to the genetic relatednes matrix, ***G***. For non-ultrametric trees, common choices of *τ* are the present time, *T*, or the mean sampling time of all the tips (Fig. II). Importantly, the specific choice of *τ* will determine the quantitative esi-mate of heritability and results in slghlty different interpretations. By choosing the present time, *τ* = *T*, the estimated heritability applies to the overall (potentially unknown) patient population that exists at the present time. Choosing *τ* as the mean sampling time across all individuals returns an estimate of heritability that applies to the pooled sampled population, represented by a hypothetical individual that was sampled at time *τ*.

## Box 3: Obtaining maximum likelihood estimates for heritability

Here, we recapitulate [33] on how to obtain heritability estimates by finding the values of (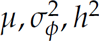) that maximize the likelihood for the SPVL data under a multivariate normal distribution with covariance matrix 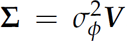. Using the expressions for the covariance matrix **Σ**(*h*^2^) from Box 1, the log-likelihood of the SPVL data, ***ϕ***, is simply

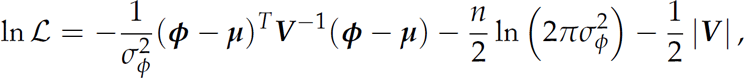

where ***μ*** = (*μ*,…, *μ*)*^T^*. The maximum likelihood (ML) estimates as a function of *h*^2^ for the mean and variance are easily found by setting the first-order derivatives of the log-likelihood equal to zero,

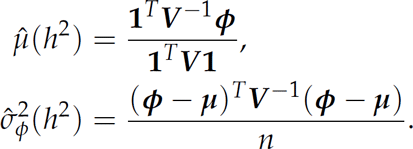

The ML estimate of *h*^2^ can then be found numerically by optimizing the log-likelihood over values of *h*^2^. Similarly, confidence intervals for *h*^2^ are derived using the profile likelihood method.

These ML estimates require computing the matrix inverse, 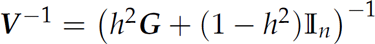 which can be computationally intensive. Historically, diagonalizing or inverting even moderately sized matrices was computationally prohibitive. Henderson [34] presented a trick that avoids matrix inversion by including unobserved ancestors as latent variables. This approach to estimating heritability is used for restricted maximum likelihood estimation in the proprietary software ASReml [35], and for Bayesian MCMC heritability estimation in the open source R package MCMCglmm [36].

However, diagonalizing even large matrices is much less prohibitive these days, allowing for a much more straightforward calculation of *h*^2^ estimates using the above expressions. Importantly, Housworth *et al.* [33] showed that estimates of *h*^2^ could be rapidly obtained by only diagonalizing the genetic relatedness matrix once, ***G*** = ***QDQ****^T^*, where ***Q*** is the matrix of eigenvectors of ***G***, and ***D*** is the diagonal matrix of eigenvalues. The normalized covariance matrix is then,

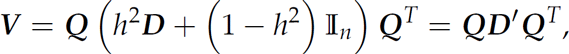

where ***D*′** is a diagonal matrix with elements 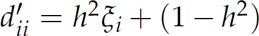, and *ξ_i_* is the *i*-th eigenvalue of ***G***. This allows for the rapid computation of the matrix inverse, ***V*^−1^ = *QD*′^−1^*Q****^T^*, where 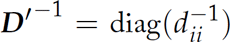. Finally, the last missing term to compute the likelihood is the determinant, 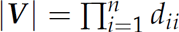.

